# Host autophagy is exploited by the intracellular parasite *Toxoplasma gondii* to enhance amino acids levels

**DOI:** 10.1101/2023.12.08.570852

**Authors:** Matthew D. White, Rajendra K. Angara, Leticia Torres Dias, Dhananjay D. Shinde, Vinai C. Thomas, Leonardo Augusto

**Affiliations:** Department of Pathology, Microbiology, and Immunology. University of Nebraska Medical Center, Omaha, NE, US; Program in Health Science, University of Santo Amaro (UNISA), São Paulo, Brazil; Cognitive Neuroscience of Development & Aging Center, University of Nebraska Medical Center, Omaha, NE, US

**Keywords:** *Toxoplasma gondii*, endoplasmic reticulum, amino acid, behavior, ER-phagy

## Abstract

*Toxoplasma gondii,* a widespread parasite, has the ability to infect nearly any nucleated cell in warm-blooded vertebrates. It is estimated that around 2 billion people globally have been infected by this pathogen. Although most healthy individuals can effectively control parasite replication, certain parasites may evade the immune response, establishing cysts in the brain that are refractory to the immune system and resistance to available drugs. For its chronic persistence in the brain, the parasite relies on host cells’ nutrients, particularly amino acids and lipids. Therefore, understanding how latent parasites persist in the brain is crucial for identifying potential drug targets against chronic forms. While shielded within parasitophorous vacuoles (PVs) or cysts, *Toxoplasma* exploits the host endoplasmic reticulum (ER) metabolism to sustains its persistence in the brain, resulting in host neurological alterations. In this study, we demonstrate that *T. gondii* disrupts the host ER homeostasis, resulting in accumulation of unfolded protein with the host ER. The host counters this stress by initiating an autophagic pathway known as ER-phagy, which breaks down unfolded proteins into amino acids, promoting their recycling. Remarkably, the persistence of latent forms in cell culture as well as behavioral changes in mice caused by the latent infection could be successfully reversed by restricting the availability of various amino acids during *T. gondi* infection. Our findings unveil the underlying mechanisms employed by *T. gondii* to exploit host ER and lysosomal pathways, enhancing nutrient levels during infection. These insights provide new strategies for the treatment of toxoplasmosis.

**Importance:** Intracellular parasites employ several mechanisms to manipulate the cellular environment, enabling them to persist in the host. *Toxoplasma gondii*, a single-celled parasite, possesses the ability to infect virtually any nucleated cell of warm-blooded vertebrates, including nearly 2 billion people worldwide. Unfortunately, existing treatments and immune responses are not entirely effective in eliminating the chronic persisting forms of the parasite. This study reveals that *T. gondii* induces the host’s autophagic pathway to boost amino acid levels in infected cells. The depletion of amino acids, in turn, influences the persistence of the parasite’s chronic forms, resulting in a reduction of neurological alterations caused by chronic infection in mice. Significantly, our investigation establishes the crucial role of host ER-phagy in the parasite’s persistence within the host during latent infection.

## Introduction

Apicomplexan parasites are among the most widespread, infecting both humans and animals on a global scale. In humans, they contribute to a spectrum of diseases with significant mortality rates, impacting billions of people worldwide. Malaria, cryptosporidiosis, and toxoplasmosis stand out as the primary diseases caused by parasites in this phylum^1^. These parasites effectively manipulate the infected cells to create the necessary niche for establishing the infection^2–5^. *Toxoplasma gondii (T. gondii)* has globally infected nearly 2 billion people due to its capability to invade virtually any nucleated cell in warm-blooded animals, leading to the development of toxoplasmosis^6^. Despite its high infection success rate, this parasite relies on host nutrients to survive. It is auxotrophic for a variety of essential components such as amino acids, lipids, and metabolites, which play a crucial role in establishing the infection^5,7–9^. To establish a persistent infection within host cells, *T. gondii* coordinates the pathways, metabolism, and organelles of infected cells to acquire nutrients^10–14^. Within the parasitophorous vacuole (PV), replicative forms (tachyzoites) have the ability to recruit infected cell organelles such as mitochondria, lysosome, and endoplasmic reticulum (ER) to acquire nutrients^9,15,16^. Notably, *T. gondii* uses the effector protein MAF1 to anchor host mitochondria, facilitating the hijacking of lipids^2,9,16^. Furthermore, *T. gondii* coordinates the protein synthesis of infected cells to enhance arginine levels to acquire this and other amino acids through the Apicomplexan Amino Acid Transporter family (ApiATs)^10,17^. In addition to utilizing amino acid transporters, *T. gondii* hijacks the host ESCRT (endosomal complexes required for transport) machinery to internalize cytosolic host proteins. These proteins are subsequently incorporated into the parasite’s vacuolar system for degradation^7,11,18,19^.

After invasion, *T. gondii* recruits the host ER to facilitate its persistence during infection^3,5^. However, the specific molecular mechanisms and key *T. gondii* proteins involved in these processes have yet to be fully identified. Surprisingly, we have previously shown that through this interaction, *T. gondii* co-opts the host ER and Unfolded Protein Response (UPR) proteins to facilitate the spread of the infection throughout the body^20^. The high-affinity association between the host ER and *T. gondii* is mediated by unknown effector protein(s), as the protein responsible for recruiting and anchoring the host ER to the parasitophorous vacuole membrane (PVM) remains unidentified. However, it has been shown that the effector protein ROP18 interacts with host ER proteins: ATF6β, MOSPD2, and RTN1-C. This interaction plays a significant role in controlling the infected cell and can contribute to the development of encephalitis in mice^3,21,22^. In addition, the secreted ROP18 phosphorylates RTN1-C in the ER, initiating ER stress-associated apoptosis through the expression of CHOP protein^22^.

The intimate association between *Toxoplasma*’s PV membrane (PVM) and the host ER induces stress in this organelle, consequently disrupting the host ER homeostasis in infected cells^20^. Given the pivotal role of ER metabolism in various cellular functions, such as autophagy, lipid synthesis, and protein folding, disruptions in ER can significantly impact overall cellular health and function^23–25^. While *T. gondii* successfully invades host cells, it lacks the machinery to synthesize specific amino acids essential for its survival and propagation. As a result, the parasite manipulates the host’s ER metabolism to acquire these crucial nutrients, effectively commandeering the host cell’s resources. This strategy approach ensures the continual existence of the parasite within the host environment. Understanding how *T. gondii* exploits host ER metabolism to access amino acids and other nutrients not only sheds light on the intricate dynamics of host-parasite interactions but also lays the foundation for potential therapeutic approaches targeting this vulnerability.

Although it has been demonstrated that *T. gondii* induces autophagy in host cells^14^, facilitating potential access to a source of amino acids and nutrients during infection, the precise molecular mechanisms driving this process are not yet fully understood. Considering the critical role of amino acid availability, especially arginine and tryptophan, in facilitating parasite replication and coordinating the parasite’s transition into the chronic stage^26,27^, our aim is to elucidate mechanisms employed by the parasite to regulate the ER metabolism of the infected cell for nutrient acquisition. Autophagic processes have been characterized to facilitate the elimination of unfolded proteins during ER stress and to maintain ER homeostasis. This mechanism, known as ER-phagy, averts the accumulation of unfolded proteins in the ER lumen, thereby preventing cell death^23,25^. During ER-phagy, the ER is sequestered within an autophagosome, which subsequently fuses with a lysosome to form an autolysosome^23,28^. Within the autolysosome, ER proteins undergo breakdown by lysosomal enzymes, leading to the release of available molecules. However, it is still unclear whether the intracellular parasite *T. gondii* exploits this process to enhance nutrient acquisition. This study addresses the mechanism by which *T. gondii* disrupts host ER homeostasis and folding capacity to elevate lysosomal amino acid levels in infected cells. We demonstrate that, throughout the course of infection, in response to the accumulation of unfolded proteins in the host ER, infected cells amplify ER-phagy. This process leads to increased levels of amino acids, including arginine, proline, and isoleucine/leucine, within the host lysosome. As a consequence, the depletion of these amino acids, which gather in the lysosomes, directly impacts the viability of chronic forms. Moreover, in the absence of amino acid intake, normal behavior is reinstated in infected mice, thereby mitigating the infection’s impact on the brain. These findings emphasize a novel role for the autophagic pathway, ER-phagy, in regulating amino acid availability during *T. gondii* infection. This mechanism plays a crucial role in enabling parasites to effectively compete with host cells for limited nutrient resources.

## Results

### *Toxoplasma gondii* infection disrupts host ER folding capacity

*T. gondii* secludes itself from the host cytoplasm within the PV forming a niche where it manipulates the host cell through secretory mechanisms, co-opting host organelles such as the host ER. This interaction results in the association of *Toxoplasma* PVM with the host ER membrane^5^. First, we confirmed the close proximity of the host ER with the *Toxoplasma* vacuole during the infection period, by utilizing an ER-Tracker in live-cell imaging to stain the ER (red). To do so, we infected Human foreskin fibroblasts (HFF) with the cystogenic strain Pru for 24h. Next, the tachyzoites were induced to differentiate into bradyzoites by incubating infected cells with alkaline media and depriving them of CO_2_. At 24-hour post infection (hpi) with tachyzoites (Tz 24h) and 4 days post-cyst formation with bradyzoites (Bz 4d), cells were incubated with the ER-Tracker for 1 h. Our data suggest that the host ER surrounds the parasite vacuole at both stages of infection (Fig. 1A).

**Figure 1.**
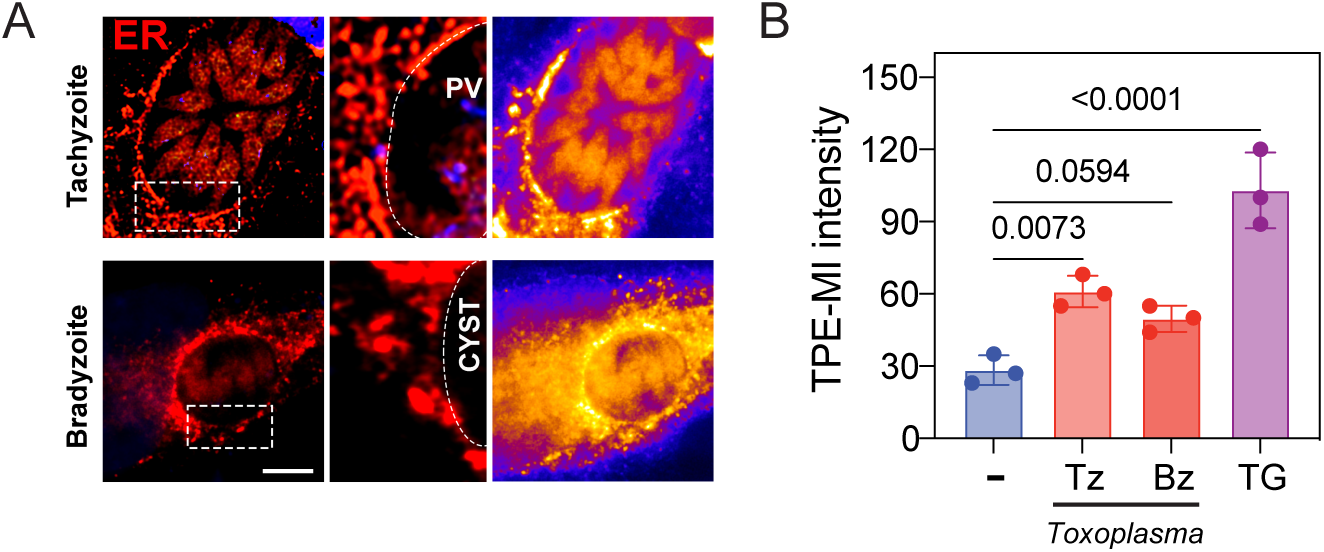
*Toxoplasma gondii* recruits host endoplasmic reticulum in both acute and chronic phases. **(A)** Representative images of tachyzoite and bradyzoite-infected cells. Infected cells were probed with ER-Tracker for live-cell imaging to stain the ER (red). Of note, ER-Tracker effectively labeled the *T. gondii* endoplasmic reticulum. The ER-Tracker is shown as a heat map with yellow showing the highest ER-Tracker intensity and blue showing the lowest ER-Tracker intensity. Scale bar = 5 μm. **(B)** Cells were incubated with TPE-MI for 10 min. TPE-MI intensity was measured in uninfected (-) and infected cells during tachyzoite (Tz) and bradyzoite (Bz) stages and normalized by the ER area using ImageJ. As a positive control, uninfected cells treated with 1 μM of Thapsigargin (TG) for 6h, an ER stress inducer. ±SD, n= 3.

Given that one of the functions of the ER is to facilitate and coordinate the folding process of proteins recently translated within the ER lumen^23,29,30^, we aimed to elucidate whether the association or close proximity between *T. gondii* and host ER dysregulates the host ER folding capacity. We previously demonstrated that *T. gondii* infection induces host UPR activation^20^. However, in this study, we were able to measure the levels of unfolded proteins in infected cells. Accordingly, we incubated infected cells with the thiol probe TPE-MI (Tetraphenylethene maleimide), which binds to cysteines of unfolded proteins and becomes fluorescent. Using live-cell microscopy, we measured the TPE-MI intensity exclusively within the ER region by transfecting cells with the ER resident protein (KDEL) tagged with RFP (red) to establish its co-localization with ER staining. This methodology enabled us to differentiate and quantify the host ER TPE-MI levels, effectively removing the contribution of parasite and cytosolic background TPE-MI intensity from our measurements. At the 24 hpi (Tz 24h) and 4 days after cyst formation (Bz 4d), cells were exposed to TPE-MI for 10 minutes. Our findings indicate a higher intensity of TPE-MI within the ER of *Toxoplasma*-infected cells compared to uninfected cells (Fig. 1B and supplementary Fig. 1). To further confirm that TPE-MI intensity was associated with the accumulation of unfolded protein and ER stress, cells were treated with the ER stress inducer thapsigargin (TG) for 6 h, revealing an even higher TPE-MI intensity in the ER (Fig. 1B and supplementary Fig. 1). Collectively, our results suggest that *T. gondii* recruits and maintain the interaction with the host ER during both the replicative and dormant stages of infection, significantly enhancing unfolded protein levels in the host ER.

### Host ER-phagy is induced in *Toxoplasma*-infected cells

Cells have evolved distinct mechanisms to alleviate the accumulation of unfolded proteins in the ER lumen through the activation of UPR^31^. Consequently, this leads to reprogramming of gene expression and translation. Remarkably, prolonged and persistent ER stress can trigger cell death. However, to prevent damage, cells can reinforce the ER stress response by activating the autophagic pathway to degrade and reduce the levels of unfolded proteins in the ER^23,29,32,33^. To investigate whether cells undergo ER-phagy in response to ER stress during *T. gondii* infection, cells were transfected with the ER resident protein (KDEL) tagged with GFP (green) and mCherry (red), which faced the cytosol. Subsequently, the cells were infected with *T. gondii*. During ER-phagy, the ER protein is targeted to autophagosomes/lysosomes, where the acidic pH destabilizes GFP, resulting in the detection of only the mCherry signal (Fig. 2A). After 30 hpi, we quantified the number of cells exhibiting punctate signals with both mCherry and GFP or only mCherry signals to determine if infected cells undergo ER-phagy. As a positive control, we used an ER-stress inducer (Thapsigargin-TG). The significant increase in mCherry signal in *Toxoplasma*-infected cells provide evidence that *T. gondii* stimulates ER-phagy in the host cells (Fig. 2B and C). Considering that ER-phagy is facilitated by FAM13B within the ER^23^, we proceeded to assess the protein levels of FAM13B in the infected cells. Our findings indicate an increase in FAM13B expression, suggesting that the infected cells undergoing ER-phagy (Fig. 2D). Together, our results suggest that *T. gondii* induces ER stress and accumulation of unfolded protein in the ER lumen. In addition to UPR activation, infected cells activate ER-phagy to alleviate the ER stress.

**Figure 2.**
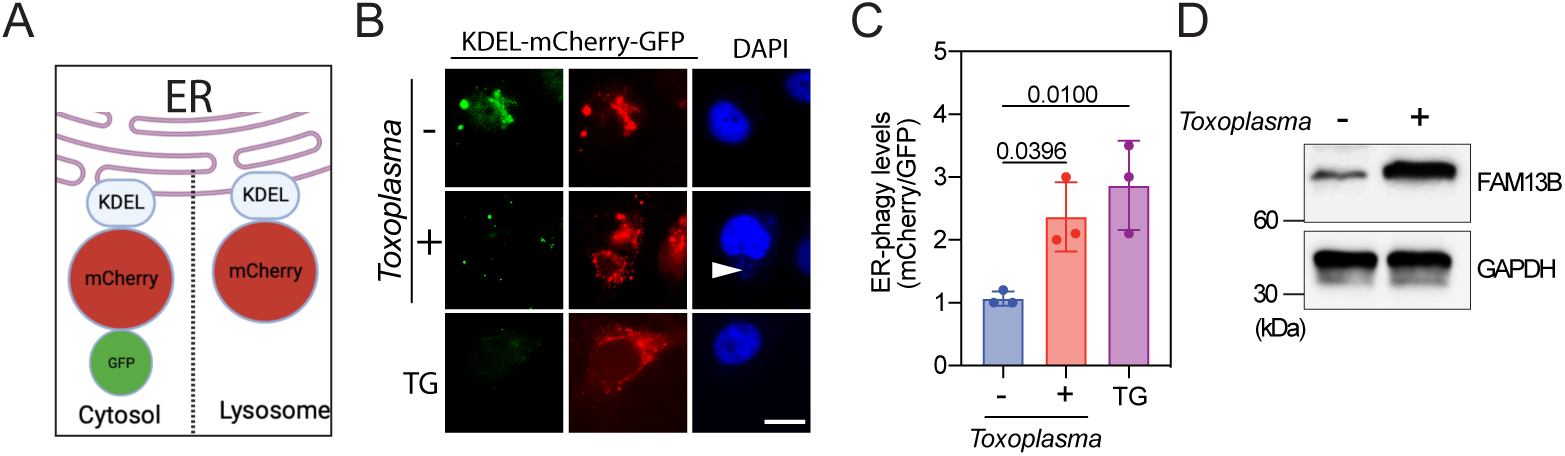
ER-phagy is induced in *Toxoplasma*-infected cells. **(A)** Schematic representation of the plasmid used to transfect cells: ER resident protein (KDEL) tagged with mCherry and GFP. Upon ER-phagy, the ER protein is integrated into lysosomes, causing instability of GFP in acidic pH, and only the mCherry signal can be detected, whereas in unstressed cells, the protein faces the cytosol, and both GFP and mCherry signals are detected. **(B-C)** Transfected cells were infected with *T. gondii* and after 30 hpi, the GFP and mCherry foci were quantified by live cell imaging of 50 cells per experiment. The data are presented as the mean ±SD, n= 3. **(D)** Then, cells were harvested and the levels of total FAM13B and GAPDH were measured by immunoblot analyses.

### Host lysosome proteolytic activity is enhanced during *Toxoplasma gondii* infection

Given that *T. gondii* infection induces ER-phagy in infected cells, we aimed to investigate whether there is an increase in proteolytic activity within the host lysosome during the infection. To accomplish this, we assessed the proteolytic activity of the lysosomal protease cathepsin B in both uninfected and infected cells, utilizing a fluorescence-based Magic Red assay. The Magic Red assay utilizes a cathepsin B peptide substrate capable of permeating the cell membrane^34^. Upon enzymatic cleavage by cathepsin B, cresyl violet generates red fluorescence, with the fluorescence intensity intensifying as the enzymatic activity progresses. Using live cell confocal microscopy, we examined lysosomal cathepsin B activity in both uninfected and *Toxoplasma*-infected cells. As our analysis revealed a higher fluorescence intensity in infected cells, and considering that cathepsin B activity is directly linked to acidic lysosomal pH^34^, our data strongly suggest elevated cathepsin B activity throughout the infection, implying a concurrent lysosomal acidification (Fig. 3).

**Figure 3.**
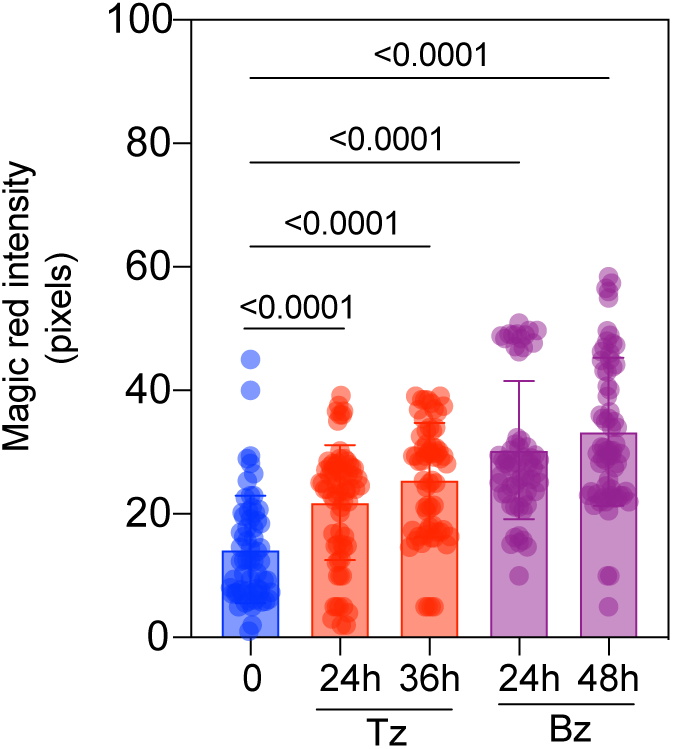
Host lysosomal proteolytic activity increases during *Toxoplasma gondii* infection. >Quantification of Magic Red reveals that lysosomes are more proteolytically active in both tachyzoite (Tz)- and bradyzoite (Bz)-infected cells compared to uninfected cells. The intensity of Magic Red (pixels) was determined by live cell imaging and quantified using ImageJ. The values were normalized to cell area and uninfected cells at each time point. Each circle represents an individual cell. The data are presented as the mean ±SD, n= 3.

### *Toxoplasma gondii* enhances lysosomal amino acid levels in the host

As ER-phagy is an autophagic process that selectively recycles ER proteins through lysosomes, and the given enhanced lysosomal proteolytic activity during infection, our next question was whether elevated host ER-phagy leads to the accumulation of amino acids in lysosomes. To investigate this, we used the LYSO-IP method, an unbiased and established technique for studying lysosomal metabolism. We generated a stable LYSO-IP cell line by inserting a copy of TMEM-192-HA (HA=hemagglutinin tag) into the genome. TMEM-192, a lysosomal protein, facilitated the rapid isolation of lysosomes from the host cytosol by HA immunoprecipitation using magnetic beads, as described^35^. Next, LYSO-IP cells were infected with *Toxoplasma*-Pru strain, and we harvested the cells at 18- and 24-h post-infection (hpi) during replicative tachyzoites (Tz) infection; or at 24- and 48-hours post bradyzoites differentiation (Bz) (Fig. 4A). Notably, we confirmed the expression of stage-specific genes for tachyzoites and bradyzoites, as well as the formation of cyst wall by identifying positive cysts using Dolichos lectin conjugated to rhodamine (Fig. 4B and C). Next, the isolation of lysosomes was confirmed by western blot using organelles specific antibodies (Fig. 4D). The lysosome fractions were collected and subjected to tandem mass spectrometry for the analysis of amino acid levels. Interestingly, we observed a time-dependent enrichment of several amino acids in lysosomes post infection when compared to mock-infected cells at each time point. Surprisingly, at 48h during bradyzoite infection (Bz 48h), the lysosomal levels of arginine, proline, lysine, isoleucine/leucine and tryptophan showed a significant increase compared to replicative tachyzoites (Fig. 4E). Importantly, all results were normalized to mock-infected cells at each time point.

**Figure 4.**
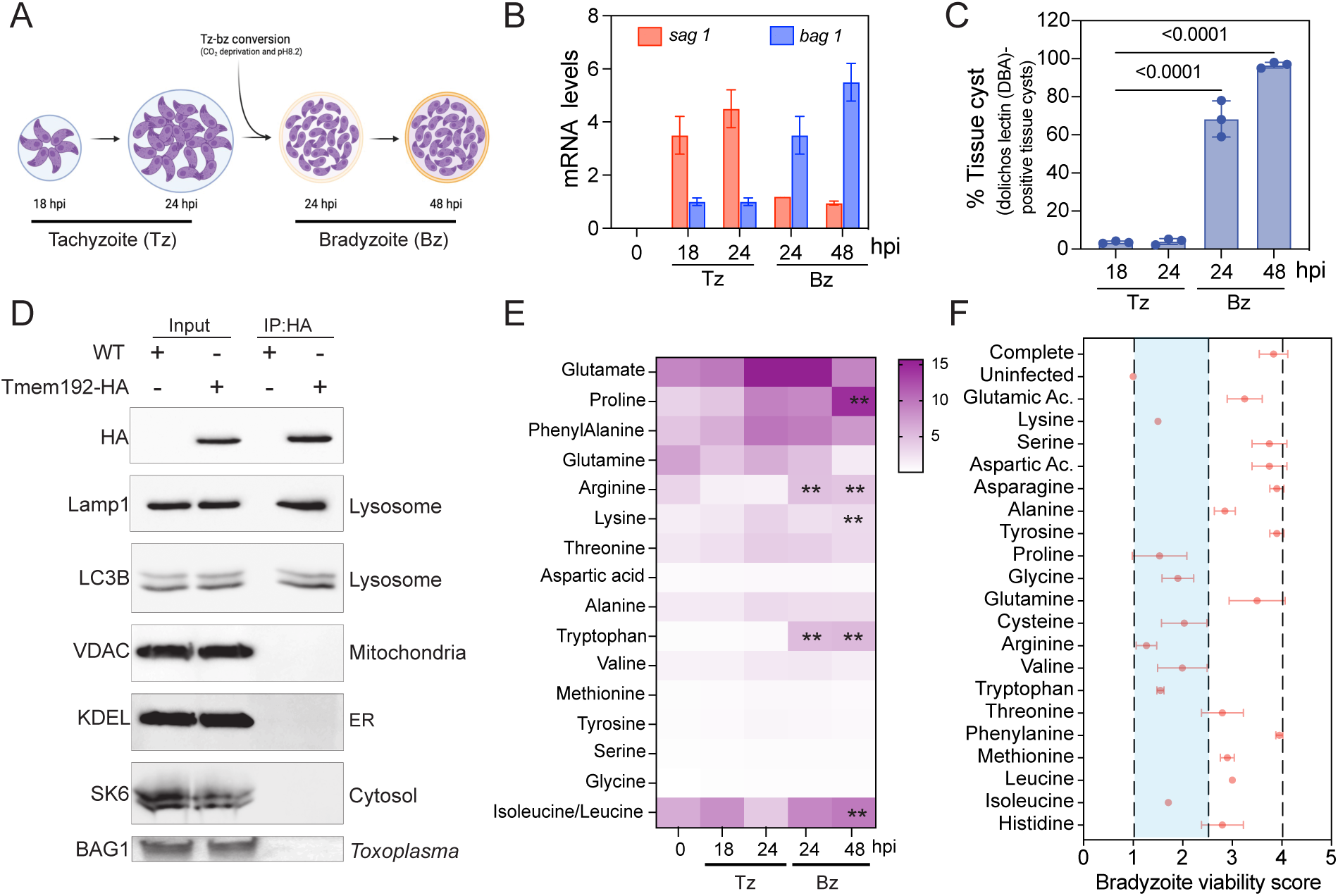
*Toxoplasma gondii* infection induces host lysosomal amino acid accumulation. **(A)** LYSO-IP cells were infected with *T. gondii* and at the indicated time points the lysosomes were isolated. **(B)** Stage-specific markers were used to confirm tachyzoite (*sag1)* and bradyzoite (*bag1)* stages during the experimental design by RT-qPCR, expression levels of both genes were normalized to *Tg-gapdh.* The basal levels were considered as 1, serving as the baseline reference for comparison. **(C)** Percentage of cyst formation was determined in 100 cells using Dolichos lectin conjugated to rhodamine to visualize cyst wall. **(D)** Host lysosomes were isolated by immunoprecipitation using HA-magnetic beads and confirmed by immunoblotting using specific organelle markers, as indicated. **(E)** Lysosomal fractions were analyzed by mass spectrometry, and the levels of amino acids were determined. The values were normalized to uninfected cells at each time point. The data are presented as the mean ±SD, n= 5. **(F)** Amino acids were individually depleted from the media and used to evaluate bradyzoite viability. After confirmation of cyst formation using Dolichos, cyst walls were lysed, and the parasites were separated from the host cell debris by filtering. Parasites were counted, and quantified by RT-qPCR. 500 parasites were used to infect a HFF monolayer. After 12 days, parasite viability was determined by plaque assay. The bradyzoite viability score was determined by the percentage of plaques number relative to completed media. The data are presented as the mean ±SD, n= 3.

Amino acids accumulated in the lysosomes due to protein degradation are transported to the cytoplasm by specific permeases^36–38^, where they can be reused in anabolic processes. Since *T. gondii* is an auxotroph for several amino acids, including arginine^26^, and the depletion of arginine significantly impairs parasite replication^39^, we have investigated the impact of availability of different amino acids on the viability of bradyzoites. To do so, cells were infected with the *Toxoplasma-*Pru strain and after cyst formation, cells were cultured for four days in a medium depleted of individual amino acids (supplemented with dialyzed fetal bovine serum (FBS)). Subsequently, the viability of bradyzoites within the cysts was assessed through pepsin digestion, followed by infecting a cell monolayer and counting the plaques. Briefly, cysts were released from infected cell layers using mechanical techniques and pepsin digestion. Parasite counts were established, and an equal number of parasites were used to infect a cell monolayer. Additionally, a portion of pepsin-treated parasites underwent genomic DNA purification for qPCR, utilizing specific primers to quantify parasite genomes per microliter. After a 12-day incubation period, plaques originating from bradyzoites were enumerated with a light microscope. The plaque count was normalized to the initial genome count, thus assessing bradyzoite viability. Bradyzoite viability in infected cells treated with the completed amino acid media was considered 100%, and the other media conditions were compared to it. Our data indicate that the depletion of arginine, proline, glycine, isoleucine, valine, lysine, tryptophan, or cysteine significantly decreases bradyzoite viability (Fig. 4F). This information underscores the critical role of amino acid availability in the context of chronic *T. gondii* infection.

### Amino acid intake levels directly affect latent toxoplasmosis *in vivo*

Given that amino acid availability has been shown to control bradyzoite viability *in vitro*, our next objective was to examine whether changes in amino acid intake during mice infection play a role in *Toxoplasma* pathogenicity. Both female and male BALB/cJ mice were intraperitoneally (i.p.) infected with type II (Pru) parasites (Fig. 5A). After 2 weeks, the infection had progressed to the chronic stage, and mice were randomized into groups. Then, we implemented custom diet formulations for up to 3 weeks (Fig. 5A). The diets were either depleted in non-essential amino acids (NoNEAA), depleted by 70% of essential amino acids (30%EAA), or amino acid complete (Normal). The mice lost less than 10% of their weight during the diet period (Fig. 5B). Additionally, no statistical difference was detected in food intake between the diets (Supplementary Fig. 2).

**Figure 5.**
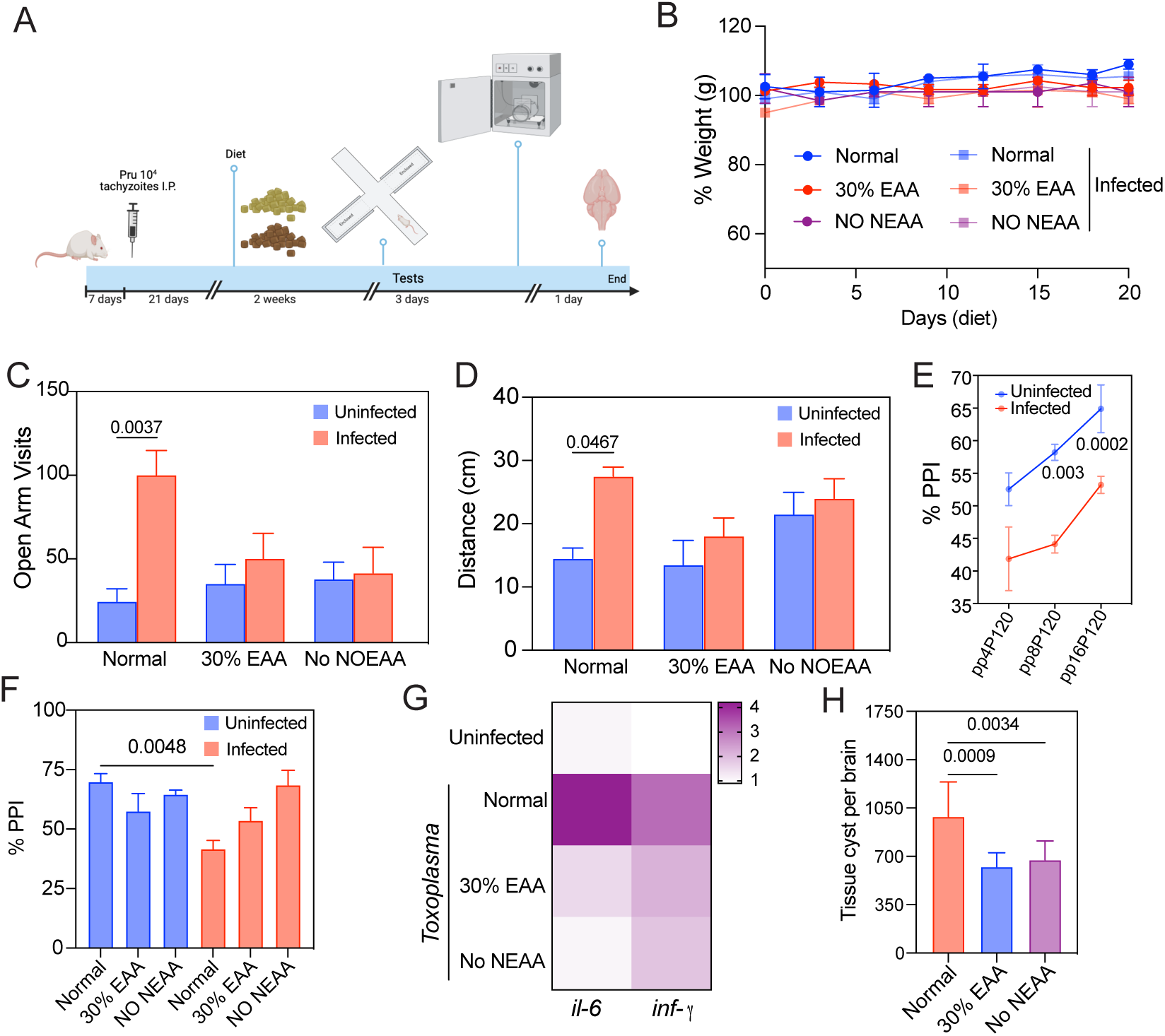
Amino acid-restricted diets partially restore behavioral changes caused by *Toxoplasma* infection in mice. **(A)** Schematic experimental protocol: BALB/cJ mice were intraperitoneally infected with 10^4^ Pru tachyzoites. After 21 days, the infected mice were randomized into groups, and amino acid-restricted diets were implemented. **(B)** Percentage of mice weight during the amino acid-restricted diet. **(C)** Quantification of visits and **(D)** distance of mice in the elevated plus maze for 60 minutes. **(E)** Percentage of PrePulse Inhibition (PPI) was determined in uninfected and infected mice. PPI is measured by the reduction in the startle response using the stimuli pp4, pp8, and pp16. **(F)** PPI was determined in uninfected and infected mice under restricted diets. PPI was measured by the reduction in the startle response using the stimulus pp16P120. **(G)** Heat map of mRNA levels of *il-6* and *inf-γ* on brains of *Toxoplasma-*infected and uninfected mice. **(H)** After the experiment endpoint, cysts were quantified in the brains of infected mice upon diet using Dolichos lectin conjugated to rhodamine in a blind manner. n= 10 female and 10 males.

As *Toxoplasma* chronic infection has been associated with behavioral changes and neurological disorders^40–43^, our initial approach involved a behavioral test - the elevated plus maze (EPM) test - to evaluate anxiety-like behavior and exploration in chronically infected mice following 2 weeks of amino acid restriction. The EPM apparatus consists of four arms arranged in a plus (+) configuration, with two arms enclosed by walls (closed arms) and two arms exposed (open arms). Mice spending more time in closed arms with fewer entries into open arms are considered to exhibit higher anxiety-like behavior, while those spending more time in open arms are seen as having lower anxiety-like behavior and greater exploration. Our results indicate that infected mice, in comparison to uninfected mice, demonstrated increased exploration and spent more time in the open arms when fed on a normal diet (Fig. 5C), suggesting reduced anxiety-like behavior. Intriguingly, custom diets lacking non-essential amino acids (NoNEAA) or containing only 30% of essential amino acids (30%EAA) restored these behavioral changes in infected mice in the EPM compared to a normal diet (Fig. 5C and D).

To deepen our understanding of the behavioral changes induced by latent *T. gondii* infection, we utilized a Prepulse inhibition (PPI) behavioral test to assess sensorimotor gating. Sensorimotor gating reflects the brain’s ability to filter out irrelevant stimuli and prevent sensory overload. PPI is quantified by the reduction in the startle response when the pre-pulse precedes the startle stimulus. PPI deficits are directly linked to various neurological impairments, including vision, hearing, memory and learning^44^. Our findings strongly indicate that *T. gondii* enhances PPI deficits during the early phase of chronic infection in mice (Fig. 5E). Surprisingly, diets lacking NoNEAA or containing 30% EAA, which were deficient in amino acids, were able to restore the PPI deficits induced by early latent *T. gondii* infection (Fig. 5F).

As previously described, behavioral changes result from the neuroinflammation caused by *T. gondii* infection^40,41^. To assess mRNA levels of *inf-γ* and *il-6* in cortex of infected mice, we utilized RT-qPCR. Our results indicated an upregulation of both genes in the brain cortex of infected mice; however, both diets, NONEAA or 30% EAA, showed a decrease in *inf-γ* and *il-6* levels (Fig. 5G). This confirmed the direct correlation between neuroinflammation and behavioral impairments caused by latent toxoplasmosis in mice. Lastly, we quantified the number of cysts present in the brain for each diet. Our findings indicate that diets lacking amino acids slightly reduce the number of cysts in the brain. However, the disease and cysts in the brain were not eliminated (Fig. 5H). Based on our data, it is strongly suggestive that the availability of amino acids plays a coordinated role in the persistence of chronic forms. Unfortunately, our limited understanding of the mechanisms and signaling pathways that *T. gondii* employs to manipulate the host’s amino acid metabolism impedes the development of innovative strategies to eliminate the infection and regulate the neurological abnormalities caused by chronic forms.

In addition, when infected mice were placed on a diet lacking all essential amino acids (No EAA), 80% of the group displayed acute seizures (Racine scale, stages 3 to 5). This manifestation could indicate a reactivation of bradyzoites in the central nervous system (CNS) (Fig. 6A). To investigate whether the seizures are linked to the reactivation of the infection, we measured the parasite burden using RT-qPCR. Specifically, we quantified the conversion of bradyzoites to tachyzoites using stage-specific primers, such as SAG1 (for tachyzoites) and BAG1 (for bradyzoites). Our findings indicate elevated levels of tachyzoite gene (SAG1) and a reduction in bradyzoite gene (BAG1), suggesting a potential reactivation of the infection due to this specific diet (Fig. 6B and C). Furthermore, the surviving mice in this group exhibited elevated *inf-γ* mRNA levels in the brain compared to infected mice on a regular diet (Fig. 6D). This suggests that the imbalanced diet intensified neuroinflammation, providing a favorable environment for the parasite. Collectively, a therapeutic approach aimed at reinstating nutrient balance and enhancing the host’s response against latent infections would be instrumental in eradicating untreated chronic forms in the brain.

**Figure 6.**
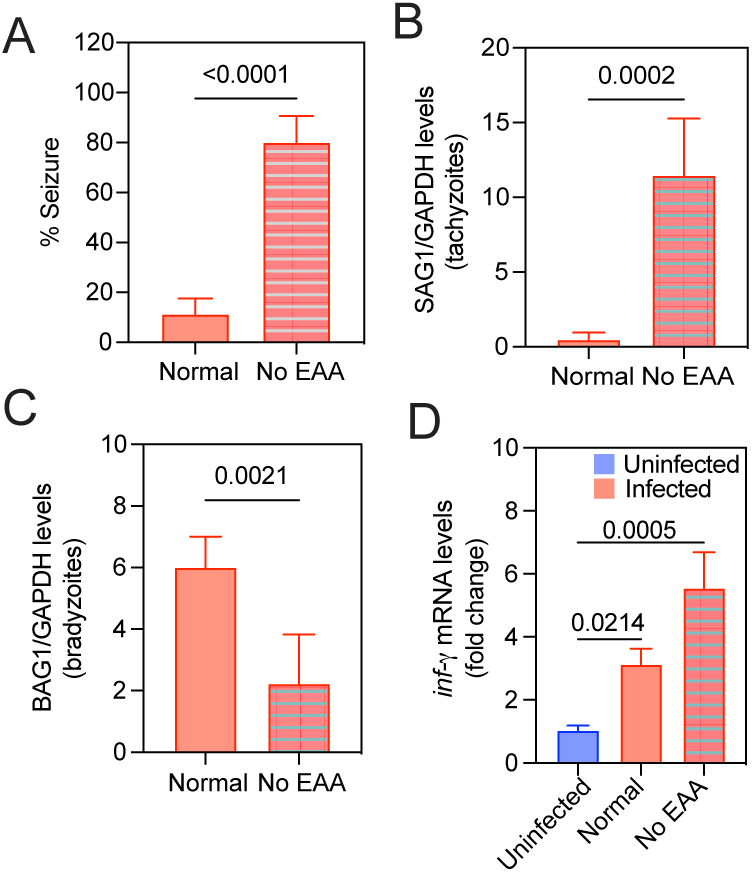
Dietary essential amino acids shape the brain environment and inflammation during *Toxoplasma gondii* infection, causing seizures. **(A)** Non-induced seizures were quantified in infected mice upon normal or essential amino acid-depleted diet using the Racine scale (stages 3 to 5). **(B-C)** Parasites levels were determined by RT-qPCR using tachyzoite- and bradyzoites- stage specific (SAG1 and BAG1) genes in the cortex of infected mice upon completed amino acid diet (normal) or a diet lacking essential amino acid (No EAA). **(D)** *inf-γ* mRNA levels were determined by RT-qPCR in surviving mice brains.

While acute *T. gondii* infection results in alterations in gut microbiota abundance^45^, such dysbiosis is not observed during the chronic phase^46^. We investigated whether dietary changes could potentially influence the modulation of gut microbiota abundance during chronic infection. Initially, we analyzed the microbiota’s at the phylum level and measured alpha diversity after two weeks of the diet in chronically infected mice, comparing it with the uninfected group. As previously reported, we did not observe any significant differences between the microbiota of chronically infected and uninfected mice under amino acid completed diet (Normal) (Fig. 7). However, under a depleted amino acid in the diet, the microbiota composition underwent a significant change, notably with an increase in Verrucomicrobia and decrease in Bacteroides. Unfortunately, our current knowledge gaps regarding the signaling pathways and the importance of brain-gut axis during chronic *T. gondii* infection and the host’s amino acid metabolism are impeding the progress of novel strategies aimed at eradicating the infection and managing the neurological complications arising from chronic forms.

**Figure 7.**
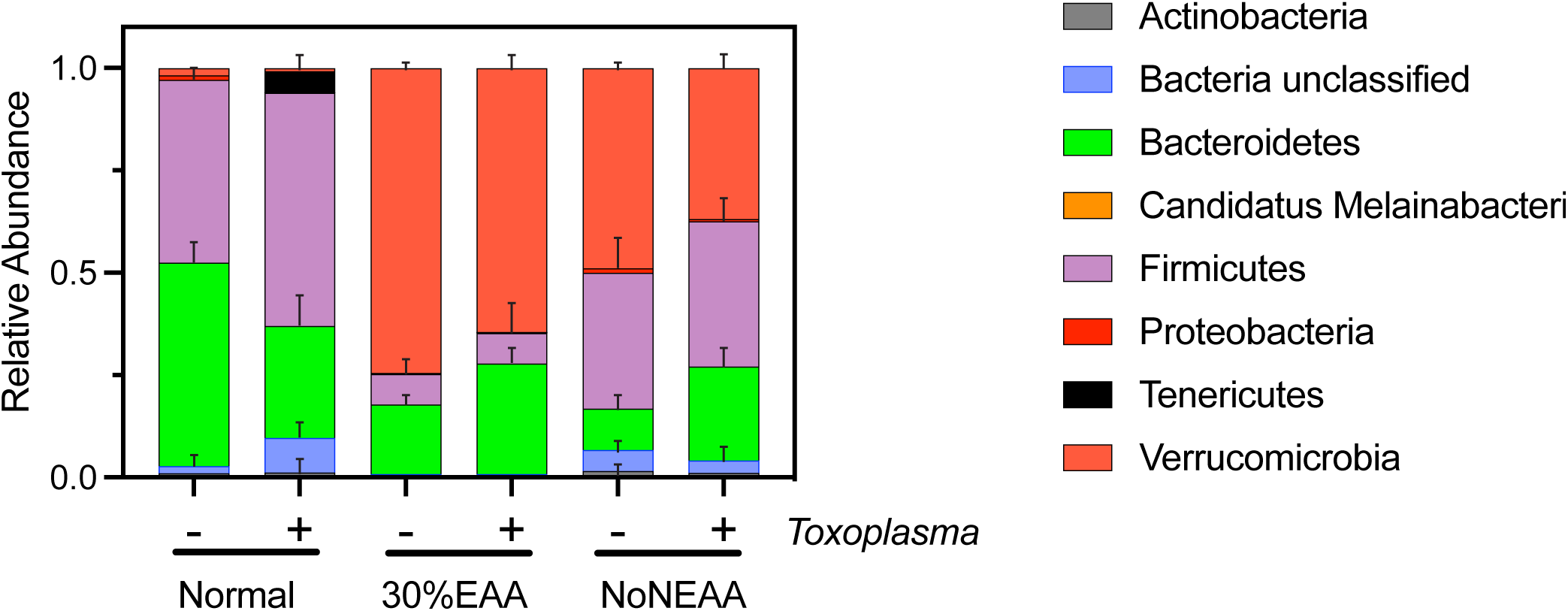
Amino acid depletion changes the microbiota composion during *Toxoplasma gondii* infection. Diversity plots depicting the composition of phyla in both uninfected and chronically infected mice under conditions of restricted amino acid intake.

## Discussion

Intracellular pathogens typically sequester themselves within vacuoles, which subsequently fuse with host cell organelles to obtain essential nutrients necessary for establishing infection^8,47^. For instance, *T. gondii* parasites inhabit a nonfusogenic parasitophorous vacuole (PV) that closely associates with host organelles, such as the endoplasmic reticulum (ER) and mitochondria. Additionally, *T. gondii* can co-opt host endosomal vesicles and cytosolic proteins to acquire nutrients^5,48–50^. The recruitment of host organelles to the PV is thought to enable *T. gondii* to modulate crucial host cell functions, including antigen presentation, nutrient production, and the suppression of apoptosis^4,5^.

In this study, we aimed to investigate the intricate mechanisms underlying the connection between the *T. gondii* vacuole and the host ER, specifically elucidating how this interaction enhances amino acid availability to support parasite survival. Our findings demonstrate a significant link between the infection and the ER, resulting in an increased presence of unfolded proteins within the ER lumen of infected cells. This, in turn, prompts infected cells to activate the UPR^20^ and initiate ER-phagy, which increased available amino acid pools. As a result, lysosomes in infected cells differentially accumulate several amino acids, potentially available for the parasite’s use. The outstanding question centers on how the parasite accesses these lysosomal amino acids. One potential mechanism involves incorporating these nutrients into the vacuole through a process similar to that observed with Rab-positive vesicles^48,51^. Alternatively, *T. gondii* may directly interact with lysosomes, inducing nutrient transport to the cytosol or causing lysosomal damage to gain access to these nutrients. Our findings highlight the importance of specific amino acids in maintaining bradyzoite viability during the chronic phase *in vitro*. However, the precise mechanism by which *T. gondii* acquires these lysosomal metabolites or enhances nutrient availability within infected cells to ensure its survival remains an elusive aspect that requires further exploration.

*T. gondii* is known to be auxotrophic for arginine and tryptophan^4,26^. However, our study emphasized the significance of amino acids such as proline, which were found to be crucial for maintaining the viability of the parasite in the cyst *in vitro*. Furthermore, in animal models, the restriction of amino acids intake has a significant impact on parasite pathogenicity. Our findings suggest that a minor reduction in the number of cysts in the brain, occurs as a result of either non-essential amino acid restriction or when only 30% of essential amino acids are present. As anticipated, these restrictions do not fully eradicate the infection; however, they lead to a reduction in inflammation, which directly correlates with changes in behavior observed in plus maze test. Upon depletion of all essential amino acids, we found strong evidence suggesting the reactivation of infection. One hypothesis is that this reactivation is due to the alleviation of inflammation or a reduction in the pressure of inflammation against the chronic forms in the brain. This creates a favorable environment that enables the parasite to sense and reactivate the infection.

Interestingly, we observed a deficiency in the PPI (Prepulse Inhibition) test during chronic infection. The PPI test can reveal alterations in memory, learning, hearing, and vision, which have been linked to *Toxoplasma* infection^40^. The precise mechanism behind these alterations remains unclear and is a topic of debate. It is uncertain whether these changes are a result from modifications in neurological function, disruptions in neuronal circuits, or inflammation caused by the cyst load in the brain regions responsible for coordinating these functions. Understanding the neurological changes during chronic infection has been challenging, primarily due to the absence of specific tropisms in the brain, as the cysts have been observed to be randomly distributed throughout the brain. An ongoing question remains as to whether the PPI deficit persists throughout the chronic infection or if it occurs only in the early stages of the chronic infection as showed in this study.

It is noteworthy that not only specific nutrients such as amino acids, but also lipids, can significantly influence *T. gondii* pathology in the brain. A crucial aspect for future research is understanding how various nutrients and metabolites collectively contribute to neuropathology during chronic infection. Therefore, understanding the metabolic dependence of bradyzoites and their host during latent infection, as well as the factors enabling parasite persistence, is crucial for the development of new therapies and the elimination of the parasite.

## Material and Methods

### Host cell and parasite culture

*Toxoplasma gondii* parasites (Prugniaud (Pru) strain-type II) were propagated in human foreskin fibroblast cells (HFF, ATCC) using Dulbecco’s modification of Eagle’s medium (DMEM) supplemented with 10% heat-inactivated fetal bovine serum (FBS) and penicillin-streptomycin. The parasites were obtained from chronically infected BALB/cJ mice. HeLa and human embryonic kidney (HEK293) cells were cultured in DMEM supplemented with 10% FBS and penicillin-streptomycin at 37°C with 5% CO_2_. Infection was performed using a multiplicity of infection (MOI) of 3 with the Pru strain for the indicated time points. The tachyzoite-bradyzoite conversion was performed by incubating infected cells with alkaline media and deprivation CO_2._

### TPE-MI staining

To prepare a stock solution of TPE-MI, TPE-MI was dissolved in DMSO at a concentration of 2 mM. HFF cells were plated in ibidi-treated channel μ-slide VI0.4 (1 × 10^4^ cells per channel; ibidi) and allowed to adhere overnight. Cells were infected with *Toxoplasma* (Pru strain) at a multiplicity of infection (MOI) of 3 for 2 hours and then washed with medium to remove extracellular parasites. After 24 hours, infected cells were incubated with freshly diluted TPE-MI (50 μM in media) for 10 min at 37 °C. The TPE-MI solution was then washed out with media, and Z-stacked confocal images were obtained for each cell using a Nikon Eclipse Ti2 spinning disk confocal microscope with a 60× oil immersion objective and an Okolab Bold Line stage top incubator. The fluorescence intensity of TPE-MI in each cell was normalized by the cell area using ImageJ software. At least 50 cells were imaged per condition for each of three independent experiments.

### Cathepsin B activity

Cathepsin B was measured using Magic Red as previously described^34^. Briefly, HFF or HeLa cells were plated in ibidi-treated channel μ-slide VI0.4 (1 × 10^4^ cells per channel; ibidi) and allowed to adhere overnight. Cells were infected with *T. gondii* (Pru strain) at a multiplicity of infection (MOI) of 3 for 2 hours and then washed with medium to remove extracellular parasites. At indicated time points, tachyzoite or bradyzoite parasites were visualized, then uninfected and infected cells were incubated with 50 μL diluted Magic Red (ImmunoChemistry Technologies) in phenol-red free DMEM and incubated for 30 min at 37°C and 5% CO_2_, and cells were live imaged, with identical capture settings, using z-stacks of 0.3-μm steps with a Nikon spinning disk confocal microscope. The Magic Red fluorescence intensity was measured, using ImageJ software, and normalized to the cell area. At least 20 cells were measured per condition in each of three independent experiments.

### ER-phagy levels

HFF or HEK293 cells (3 × 10^4^ cells per well of a 24-well plate or 1 × 10^5^ cells per well of a 6-well plate) were reverse transfected with 0.5 μg of pCW57-CMV-KDEL-mCherry-GF using Fugene HD (Promega) according to the manufacturer’s instructions. After 24 h post-transfection, cells were replated in ibidi-treated channel μ-slide VI0.4 (1 × 10^4^ cells per channel; ibidi), and infected with *T. gondii* for 2 h (MOI, 3). Cells were live imaged, with identical capture settings, using z-stacks of 0.3-μm steps with a Nikon spinning disk confocal microscope. At least 25 cells were measured per condition in each of three independent experiments.

### Immunoblot

Infected cells were harvested in RIPA buffer solution supplemented with cOmplete, EDTA-free protease inhibitor cocktail (Roche). Protein quantification was performed using the BCA Protein Assay Kit (Pierce). Equal amounts of protein lysates were separated by 4–20% Mini-PROTEAN TGX (BioRad), and proteins were transferred to a nitrocellulose membrane. Immunoblot analyses were done using primary antibodies (Cell signaling: FAM13B, GAPDH, HA, LAMP1, LC3B, VDAC, KDEL, SK6) for 18h, followed by secondary antibody horseradish peroxidase (HRP)-conjugated. Proteins in the immunoblot membranes were visualized using the Azure Biosystem C600. Immunoblot analyses were carried out for three independent experiments.

### Lysosome immunoprecipitation

HEK293-TMEM192-HA cells were plated (3 × 10^6^ cells per 150-mm tissue-culture dishes). After 24h, cells were infected with *Toxoplasma*-Pru for 2 h (MOI, 3) and then harvested at the indicated time points. The cyst formation was induced by alkaline stress (pH 8.2) combined with CO_2_ deprivation. At the indicated time points, cells were washed three times with cold KPBS (136 mM KCl, 10 mM KH_2_PO_4_, pH 7.3), scraped on ice in 1 mL of cold KPBS and centrifuged at 900 x g for 2 min at 4°C. The pellets were resuspended in 950 μL and 25% of each sample was reserved as whole-cell fraction, for further processing by LC/MS/MS analysis. The remaining cells were gently homogenized with 15 strokes of a 2 ml homogenizer, the lysates were then centrifuged at 900 x g for 2 min at 4°C. Next, the supernatant containing the lysosomes was incubated with KPBS prewashed anti-HA magnetic beads (Pierce) for 5 min. Isolated lysosomes were then gently washed five times with KPBS using the DynaMag Spin Magnet. The amino acid and metabolite extraction from lysosomes was performed by incubating the isolated lysosomes with 100 μL of metabolite methanol extraction buffer (80% methanol, 20% water containing internal standards) for 10 min on ice, followed by beads removal. The metabolite extract (liquid fraction) was then centrifuged at 900 x g for 5 min at 4°C. The supernatant was collected and analyzed by LC-MS/MS to determine the amino acid levels. Mass spectrometry analysis was performed by the Mass-Spectrometry and Proteomics Core Facility at University of Nebraska Medical Center. 13C15N-labeled canonical amino acid (CAA) mix procured from the Cambridge Isotope Laboratory was used as the internal standard during the sample preparation.

### LC-HRMS/MS analysis of Metabolites

A high-resolution mass spectrometer, specifically the Tribid Orbitrap Exploris 480 (Thermo) connected to an ultra-high-performance liquid chromatography (UHPLC) system, was employed for metabolite analysis. The chromatographic separation was performed by liquid chromatography using XBridge Amide (150mm × 2.1mm ID; 1.7µm particle size) analytical column and a binary solvent system with a flow rate of 0.3 ml/min. Mobile phase A was composed of 10 mM ammonium acetate and 10 mM ammonium hydroxide containing 5% acetonitrile in LC-MS grade water. Mobile phase B was 100% LC-MS grade acetonitrile. The column temperature was maintained at 40°C, while the autosampler was set to 5°C. UHPLC pumps operated in gradient mode, and 5 µL injection volume was used per sample.

For untargeted metabolomics in data-dependent MS/MS acquisition mode (DDA), the HRMS Orbitrap (Exploris 480) was employed in polarity switching mode. Electrospray ionization (ESI) parameters were optimized with −3500V and 4000V electrospray ion voltage in negative and positive modes, respectively. The ion transfer tube temperature was set to 400°C, and the m/z scan range was 70-1050 Da. Sheath gas, auxiliary gas and sweep gas were optimized according to the UHPLC flow rate. Orbitrap resolution for precursor ion as well as for fragment ion scan was maintained at 240000 and 120000 respectively. Normalized collision energies at 30, 50 and 150% were used for the fragmentation. Data acquisition was performed in profile mode using Xcaliber software (Thermo). The software includes Qual-, Quant-, and FreeStyle browsers, which were utilized for profiling labeled 13C15N-CAA standards in all samples. Identification and detection of metabolites were supported by Compound Discoverer (CD) software (Thermo). The HMDB and KEGG databases were integrated for metabolite identifications. The high-resolution mass spectrometry allowed for accurate precursor and fragment ion mass detection, facilitating confident molecular annotation and metabolite assignments.

### Bradyzoite viability

After cyst formation, the cells were cultured for four days in RPMI medium with depleted individual amino acids, supplemented with dialyzed fetal bovine serum (FBS). The depleted medium was prepared by adding 19 amino acids to the RPMI medium lacking one amino acid. After incubation, cysts were released from infected cells using mechanical techniques and pepsin digestion. Then, parasite counts were determined, and an equal number of parasites (500) was used to infect an HFF monolayer. Moreover, a subset of pepsin-treated parasites underwent genomic DNA purification for qPCR, using specific primers (B1)^20^ to quantify the number of parasite genomes per microliter. Following a 12-day incubation period, plaques were counted using a light microscope. The plaque count was standardized according to the initial genome and parasite counts, thus evaluating the viability of bradyzoites. Bradyzoite viability in infected cells, treated with the complete amino acid medium, was set as 100%, and the other medium conditions were compared against it. Each experiment was performed three times, each with three technical replicates.

### Mice infection

The mice used in this study were housed in an animal facility-UNMC Animal Care Comparative Medicine at University of Nebraska Medical Center. The Institutional Animal Care and Use Committee (IACUC) at the Office of Vice Chancellor for Research Comparative Medicine approved the use of all animals and procedures (IACUC protocol number 2106411).

BALB/cJ mice were purchased from Jackson Laboratory at 6 weeks of age. After one week of acclimation, mice were intraperitoneally infected with 10^4^ Pru tachyzoites. Mock-infected animals were handled identically and injected with the same volume of sterile PBS. After 21 days post-infection, mice were randomized into groups. Blood samples were extracted to confirm parasite infection by serological analysis using a dot blot containing parasite lysate. Then, three different diets were implemented for up to 2 weeks. The mice weight and seizure were measured three times per week. Each group had 10 males and 10 females.

To assess bradyzoite reactivation, DNA was extracted from the brain and quantified using qPCR to measure the copy number of stage-specific genes for tachyzoites or bradyzoites. SAG1-F: GCTGTAACATTGAGCTCCTTGASTTCCTG; R: CCGGAACAGTACTGATTGTTGTCTTGAG. BAG1-F: AGTCGACAACGGAGCCATCGTTATC; R: ACCTTGATCGTGACACGTAGAACGC.

### Behavioral and cognitive analyses

After implementation of diets, mice were subjected to the behavioral analyses: elevated plus-maze (EPM), and prepulse inhibition (PPI). The mice were transferred to the Animal Behavior Core and the technical team operated the experiments. For the elevated plus-maze (EPM), mice were placed in the center of the apparatus, facing a closed arm, and each mouse was recorded for 60 minutes. The time and distance spent in the open and closed arms were measured by video and analyzed by software. Finally, the Prepulse inhibition of the startle test was performed with pulse-alone trials using 120 dB of white noise. Subsequently, blocks of trials were presented. Each block consisted of one pulse-alone trial and three prepulse-alone trials (+4, +8, or +16 units). The startle point was measured in milliseconds. All the experiments were performed in a blinded manner.

### Seizure scoring

Seizure monitoring and scoring were performed as described^52,53^. Briefly, mice were observed daily and graded using the Racine scale. Stage 1: mouth and facial disturbances; Stage 2: head nodding; Stage 3: forelimb clonus; Stage 4, rearing; Stage 5: falling^52,53^. The percentage of mice with seizures was calculated by the number of mice with seizures/total number of mice per group × 100.

### Measurement of mRNA levels

Total RNA of cortex lysate was isolated from the cells using Trizol LS (Invitrogen), and cDNA was generated using Omniscript (Qiagen) as described by the manufacturer. RT-qPCR was carried out using SYBR Green real-time PCR master mixes (Invitrogen) and the StepOnePlus Real system (Applied Biosystems). Relative levels of transcripts were calculated with the *ΔΔCT* method. Each experiment was performed blinded three times, each with three technical replicates. Primers: IL-6: forward-TTCCATCCAGTTGCCTTCTT, reverse: TCCACGATTTCCCAGAGAAC; GAPDH forward: TGCACCACCAACTGCTTAG, reverse: GGATGCAGGGATGATGTTC; IFN-γ forward: TTCTTCAGCAACAGCAAGGC, reverse: TCAGCAGCGACTCCTTTTCC.

### Quantification and statistical analysis

Quantitative data are presented as means and standard deviations and were derived from at least three biological replicates. Statistical significance was determined using one-way analysis of variance (ANOVA) with Tukey’s post hoc test and multiple two-tailed t test using Graph Prism software 10. The number of biological replicates and P values are indicated in figure legends. For immunoblot analyses, the reported images are representative of at least three independent experiments.

### 16S rRNA Sequencing and Analysis

Fecal pellets were collected for individual mouse in each group at the same day and stored in DNA/RNA shield solution (Zymo Reasearch) at −20°C until further processing. DNA was extracted using the QIAmp Fast DNA Stool mini kit. The samples were shipped to Zymo Research Corp for sequencing and Metagenomics analysis including alpha-diversity (richness, Shannon diversity).

## Acknowledgements

We thank Anna Dunaevsky, Stacey Gilk, and committee members of the Cognitive Neuroscience of Development & Aging Center (CoNDA) for helpful discussions and critical feedback. We thank Emily Heaton for the critical feedback on the manuscript. This research was supported by the Research Project Leader project at the CoNDA center through the Centers for Biomedical Research Excellence (COBRE) Program (NIH 1P20GM130447-01A1) to L.A., and the NIH/NIAID P01A1083211 (Metabolomics Core) to V.C.T.. We thank the Animal Behavior Core at the University of Nebraska Medical Center for conducting the behavioral tests and suggestions, the core is supported by state funds from the Nebraska Research Initiative (NRI). The University of Nebraska Medical Center Mass Spectrometry and Proteomics Core Facility is administrated through the Office of the Vice Chancellor for Research and supported by state funds from the NRI.

We have no conflicts of interest to declare.

## Author contributions

MDW, RKA, LTD, VCT, DDS and LA designed the experiments. MDW, RKA, LTD, DDS and LA performed experiments. MDW and LA wrote the manuscript.

---

**Supplementary Figure 1.**
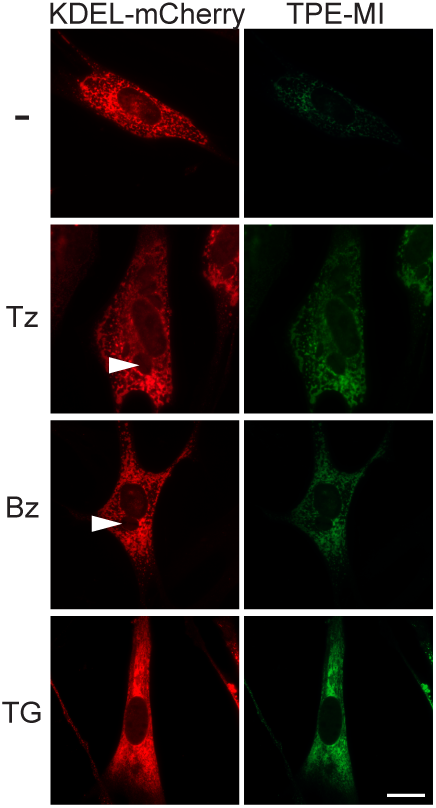
Live cell image of cells treated with TPE-MI. Cells were incubated with TPE-MI for 10 min. The intensity of TPE-MI was determined by live cell imaging and quantified using ImageJ. The positive control consisted of uninfected cells treated with 1μM Thapsigargin (TG) for 6h, an ER stress inducer. Scale bar = 5 μm

**Supplementary Figure 2.**
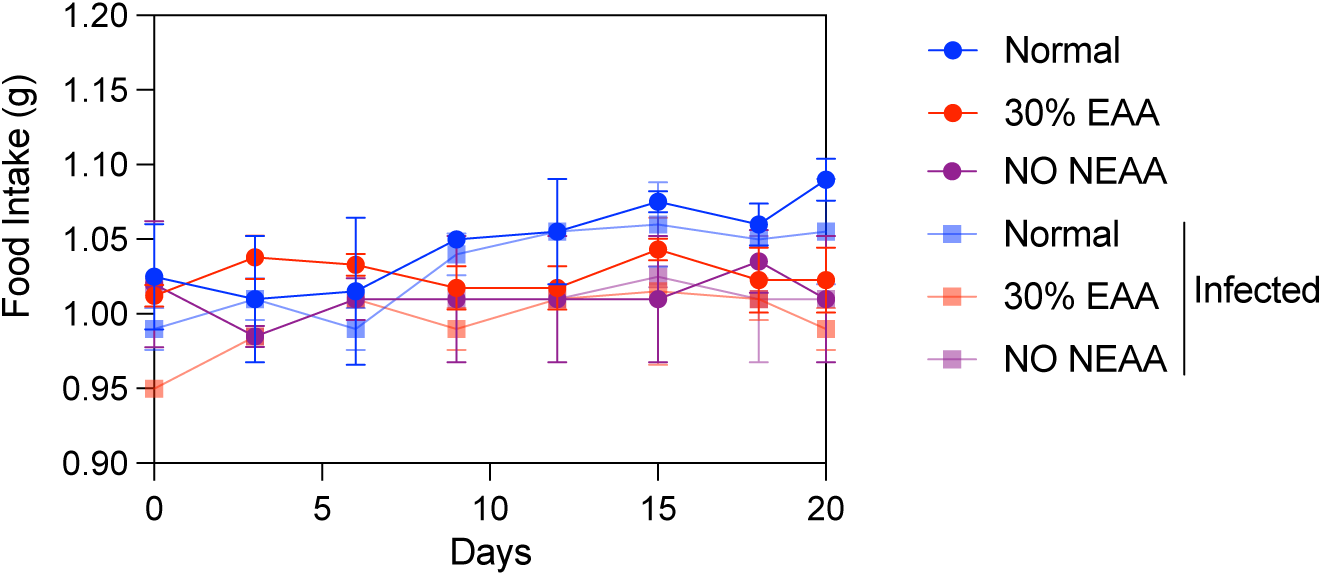
Food intake in chronic infected mice. Assessment of the proportion of food consumed by mice, both uninfected and infected, over a two-week period using a customized diet.

## Notes

### Competing Interest Statement

The authors have declared no competing interest.

## Reference

1 Delgado, I. L. S. et al. The Apicomplexan Parasite Toxoplasma gondii. Encyclopedia 2, 189–211 (2022).

2 Li, X. et al. Mitochondria shed their outer membrane in response to infection-induced stress. Science 375, eabi4343 (2022). 10.1126/science.abi4343

3 Ferrel, A., Romano, J., Panas, M. W., Coppens, I. & Boothroyd, J. C. Host MOSPD2 enrichment at the parasitophorous vacuole membrane varies between *Toxoplasma* strains and involves complex interactions. bioRxiv, 2022.2012.2023.521848 (2022). 10.1101/2022.12.23.521848

4 Augusto, L., Wek, R. C. & Sullivan, W. J., Jr. Host sensing and signal transduction during Toxoplasma stage conversion. Mol Microbiol 115, 839–848 (2021). 10.1111/mmi.14634

5 Coppens, I. & Romano, J. D. Hostile intruder: Toxoplasma holds host organelles captive. PLoS Pathog 14, e1006893 (2018). 10.1371/journal.ppat.1006893

6 Montoya, J. G. & Liesenfeld, O. Toxoplasmosis. Lancet 363, 1965–1976 (2004). 10.1016/s0140-6736(04)16412-x

7 Di Cristina, M. et al. Toxoplasma depends on lysosomal consumption of autophagosomes for persistent infection. Nat Microbiol 2, 17096 (2017). 10.1038/nmicrobiol.2017.96

8 Pernas, L. Cellular metabolism in the defense against microbes. J Cell Sci 134 (2021). 10.1242/jcs.252023

9 Pernas, L., Bean, C., Boothroyd, J. C. & Scorrano, L. Mitochondria Restrict Growth of the Intracellular Parasite Toxoplasma gondii by Limiting Its Uptake of Fatty Acids. Cell Metab 27, 886–897.e884 (2018). 10.1016/j.cmet.2018.02.018

10 Piro, F. et al. A Toxoplasma gondii putative arginine transporter localizes to the plant-like vacuolar compartment and controls parasite extracellular survival and stage differentiation. bioRxiv (2023). 10.1101/2023.08.31.555807

11 Rivera-Cuevas, Y. & Carruthers, V. B. The multifaceted interactions between pathogens and host ESCRT machinery. PLoS Pathog 19, e1011344 (2023). 10.1371/journal.ppat.1011344

12 Missiaen, R. et al. GCN2 inhibition sensitizes arginine-deprived hepatocellular carcinoma cells to senolytic treatment. Cell Metab 34, 1151–1167.e1157 (2022). 10.1016/j.cmet.2022.06.010

13 Olson, W. J. et al. Dual metabolomic profiling uncovers Toxoplasma manipulation of the host metabolome and the discovery of a novel parasite metabolic capability. PLoS Pathog 16, e1008432 (2020). 10.1371/journal.ppat.1008432

14 Wang, Y., Weiss, L. M. & Orlofsky, A. Host cell autophagy is induced by Toxoplasma gondii and contributes to parasite growth. J Biol Chem 284, 1694–1701 (2009). 10.1074/jbc.M807890200

15 Franco, M. et al. A Novel Secreted Protein, MYR1, Is Central to Toxoplasma’s Manipulation of Host Cells. mBio 7, e02231–02215 (2016). 10.1128/mBio.02231-15

16 Pernas, L. et al. Toxoplasma effector MAF1 mediates recruitment of host mitochondria and impacts the host response. PLoS Biol 12, e1001845 (2014). 10.1371/journal.pbio.1001845

17 Fairweather, S. J. et al. Coordinated action of multiple transporters in the acquisition of essential cationic amino acids by the intracellular parasite Toxoplasma gondii. PLoS Pathog 17, e1009835 (2021). 10.1371/journal.ppat.1009835

18 Rivera-Cuevas, Y. et al. Toxoplasma gondii exploits the host ESCRT machinery for parasite uptake of host cytosolic proteins. PLoS Pathog 17, e1010138 (2021). 10.1371/journal.ppat.1010138

19 Smith, D. et al. Toxoplasma TgATG9 is critical for autophagy and long-term persistence in tissue cysts. Elife 10 (2021). 10.7554/eLife.59384

20 Augusto, L. et al. Toxoplasma gondii Co-opts the Unfolded Protein Response To Enhance Migration and Dissemination of Infected Host Cells. mBio 11 (2020). 10.1128/mBio.00915-20

21 Yamamoto, M. et al. ATF6beta is a host cellular target of the Toxoplasma gondii virulence factor ROP18. J Exp Med 208, 1533–1546 (2011). 10.1084/jem.20101660

22 An, R. et al. Encephalitis is mediated by ROP18 of Toxoplasma gondii, a severe pathogen in AIDS patients. Proc Natl Acad Sci U S A 115, E5344–e5352 (2018). 10.1073/pnas.1801118115

23 Mochida, K. & Nakatogawa, H. ER-phagy: selective autophagy of the endoplasmic reticulum. EMBO Rep 23, e55192 (2022). 10.15252/embr.202255192

24 Moncan, M. et al. Regulation of lipid metabolism by the unfolded protein response. J Cell Mol Med 25, 1359–1370 (2021). 10.1111/jcmm.16255

25 Chino, H. & Mizushima, N. ER-Phagy: Quality Control and Turnover of Endoplasmic Reticulum. Trends Cell Biol 30, 384–398 (2020). 10.1016/j.tcb.2020.02.001

26 Fox, B. A., Gigley, J. P. & Bzik, D. J. Toxoplasma gondii lacks the enzymes required for de novo arginine biosynthesis and arginine starvation triggers cyst formation. Int J Parasitol 34, 323–331 (2004). 10.1016/j.ijpara.2003.12.001

27 Pfefferkorn, E. R. Interferon gamma blocks the growth of Toxoplasma gondii in human fibroblasts by inducing the host cells to degrade tryptophan. Proc Natl Acad Sci U S A 81, 908–912 (1984). 10.1073/pnas.81.3.908

28 Root, J., Merino, P., Nuckols, A., Johnson, M. & Kukar, T. Lysosome dysfunction as a cause of neurodegenerative diseases: Lessons from frontotemporal dementia and amyotrophic lateral sclerosis. Neurobiol Dis 154, 105360 (2021). 10.1016/j.nbd.2021.105360

29 Hetz, C. & Papa, F. R. The Unfolded Protein Response and Cell Fate Control. Mol Cell 69, 169–181 (2018). 10.1016/j.molcel.2017.06.017

30 Chang, T. K. et al. Coordination between Two Branches of the Unfolded Protein Response Determines Apoptotic Cell Fate. Mol Cell 71, 629–636.e625 (2018). 10.1016/j.molcel.2018.06.038

31 Walter, P. & Ron, D. The unfolded protein response: from stress pathway to homeostatic regulation. Science 334, 1081–1086 (2011). 10.1126/science.1209038

32 Hetz, C. & Saxena, S. ER stress and the unfolded protein response in neurodegeneration. Nat Rev Neurol 13, 477–491 (2017). 10.1038/nrneurol.2017.99

33 Wu, H., Ng, B. S. & Thibault, G. Endoplasmic reticulum stress response in yeast and humans. Biosci Rep 34 (2014). 10.1042/bsr20140058

34 Clemente, T. M. et al. Coxiella burnetii Sterol-Modifying Protein Stmp1 Regulates Cholesterol in the Intracellular Niche. mBio 13, e0307321 (2022). 10.1128/mbio.03073-21

35 Abu-Remaileh, M. et al. Lysosomal metabolomics reveals V-ATPase- and mTOR- dependent regulation of amino acid efflux from lysosomes. Science 358, 807–813 (2017). 10.1126/science.aan6298

36 Jézégou, A. et al. Heptahelical protein PQLC2 is a lysosomal cationic amino acid exporter underlying the action of cysteamine in cystinosis therapy. Proc Natl Acad Sci U S A 109, E3434–3443 (2012). 10.1073/pnas.1211198109

37 Liu, B., Du, H., Rutkowski, R., Gartner, A. & Wang, X. LAAT-1 is the lysosomal lysine/arginine transporter that maintains amino acid homeostasis. Science 337, 351–354 (2012). 10.1126/science.1220281

38 Sagné, C. et al. Identification and characterization of a lysosomal transporter for small neutral amino acids. Proc Natl Acad Sci U S A 98, 7206–7211 (2001). 10.1073/pnas.121183498

39 Augusto, L., Amin, P. H., Wek, R. C. & Sullivan, W. J., Jr. Regulation of arginine transport by GCN2 eIF2 kinase is important for replication of the intracellular parasite Toxoplasma gondii. PLoS Pathog 15, e1007746 (2019). 10.1371/journal.ppat.1007746

40 Boillat, M. et al. Neuroinflammation-Associated Aspecific Manipulation of Mouse Predator Fear by Toxoplasma gondii. Cell Rep 30, 320–334.e326 (2020). 10.1016/j.celrep.2019.12.019

41 Martynowicz, J., Augusto, L., Wek, R. C., Boehm, S. L., 2nd & Sullivan, W. J., Jr. Guanabenz Reverses a Key Behavioral Change Caused by Latent Toxoplasmosis in Mice by Reducing Neuroinflammation. mBio 10 (2019). 10.1128/mBio.00381-19

42 Parlog, A. et al. Chronic murine toxoplasmosis is defined by subtle changes in neuronal connectivity. Dis Model Mech 7, 459–469 (2014). 10.1242/dmm.014183

43 Hermes, G. et al. Neurological and behavioral abnormalities, ventricular dilatation, altered cellular functions, inflammation, and neuronal injury in brains of mice due to common, persistent, parasitic infection. J Neuroinflammation 5, 48 (2008). 10.1186/1742-2094-5-48

44 Schulz, S. E., Luszawski, M., Hannah, K. E. & Stevenson, R. A. Sensory Gating in Neurodevelopmental Disorders: A Scoping Review. Res Child Adolesc Psychopathol 51, 1005–1019 (2023). 10.1007/s10802-023-01058-9

45 French, T. et al. Persisting Microbiota and Neuronal Imbalance Following T. gondii Infection Reliant on the Infection Route. Front Immunol 13, 920658 (2022). 10.3389/fimmu.2022.920658

46 Saraav, I. et al. Chronic Toxoplasma gondii infection enhances susceptibility to colitis. Proc Natl Acad Sci U S A 118 (2021). 10.1073/pnas.2106730118

47 Sachdeva, K. & Sundaramurthy, V. The Interplay of Host Lysosomes and Intracellular Pathogens. Front Cell Infect Microbiol 10, 595502 (2020). 10.3389/fcimb.2020.595502

48 Hartman, E. J., Asady, B., Romano, J. D. & Coppens, I. The Rab11-family interacting proteins reveal selective interaction of mammalian recycling endosomes with the Toxoplasma parasitophorous vacuole in a Rab11- and Arf6-dependent manner. Mol Biol Cell 33, ar34 (2022). 10.1091/mbc.E21-06-0284

49 Romano, J. D. et al. The parasite Toxoplasma sequesters diverse Rab host vesicles within an intravacuolar network. J Cell Biol 216, 4235–4254 (2017). 10.1083/jcb.201701108

50 Nolan, S. J., Romano, J. D. & Coppens, I. Host lipid droplets: An important source of lipids salvaged by the intracellular parasite Toxoplasma gondii. PLoS Pathog 13, e1006362 (2017). 10.1371/journal.ppat.1006362

51 Coppens, I. et al. Toxoplasma gondii sequesters lysosomes from mammalian hosts in the vacuolar space. Cell 125, 261–274 (2006). 10.1016/j.cell.2006.01.056

52 Van Erum, J., Van Dam, D. & De Deyn, P. P. PTZ-induced seizures in mice require a revised Racine scale. Epilepsy Behav 95, 51–55 (2019). 10.1016/j.yebeh.2019.02.029

53 Racine, R. J. Modification of seizure activity by electrical stimulation. II. Motor seizure. Electroencephalogr Clin Neurophysiol 32, 281–294 (1972). 10.1016/0013-4694(72)90177-0

